# Unravelling genetic differentiation between *Glossina brevipalpis* populations from two distant National Parks in Mozambique

**DOI:** 10.1101/2025.02.28.640735

**Authors:** Denise R. A. Brito, Adeline Ségard, Fernando C. Mulandane, Nióbio Cossa, Hermógenes N. Mucache, Sophie Ravel, Thierry De Meeûs, Luis Neves

## Abstract

African trypanosomosis (AT), caused by protozoan parasites of the genus *Trypanosoma*, has plagued the African continent for centuries, affecting both humans and animals. Vector control represents an efficient way to reduce the burden of AT. In Mozambique, control campaigns reshaped tsetse fly distribution to what it is today, with *Glossina brevipalpis* and *G. austeni* as the dominant species in the South. Additionally, *G. brevipalpis* can be found in two National parks, Gorongosa National Park in the Centre and Maputo National Park in the South, with an 840 km wide tsetse-free zone between them. In order to improve our knowledge of the genetic diversity and gene flow of these populations and their probable isolation, we undertook a population genetics study with 11 microsatellite loci. We found that these two zones behave as isolated subpopulations, only exchanging a few individuals per year. To explain this finding, we suggest the existence of undocumented pocket populations between the two parks, or, in the absence of these, the accidental translocation of tsetse flies during human-driven animal transportation. We concluded that such undocumented means of dispersal should be explored further in studies investigating *Glossina* populations, to allow for the design of more efficient control strategies.

## Introduction

African trypanosomosis (AT), caused by protozoan parasites of the genus *Trypanosoma*, has plagued the African continent for centuries, affecting both humans and animals, causing a heavy economic burden [1–4]. These blood parasites have a diverse range of mammalian hosts [1]. Their primary biological vectors are the hematophagous flies known as tsetse (*Glossina* spp). Control of tsetse flies has been conducted over the last and present century so as to reduce the burden caused by both human (HAT) and animal (AAT) African trypanosomosis [3,5,6].

*Glossina* species are classified into three main groups (*Fusca*, *Palpalis* and *Morsitans*). These presently include a total of 31 species/subspecies adapted to specific habitats [3,5,7]. The *Fusca* group corresponds to forest-dwelling flies that are mostly found in western-central Africa, except for *Glossina brevipalpis* and *G. longipennis,* which are located in eastern and southern Africa [3,5]. *Glossina brevipalpis* is characterised as a large fly of the *Fusca* Group [3,5,8]. According to data published in the last 30 years, *G. brevipalpis* populations have been detected in Kenya, Rwanda, Tanzania, Zambia, Mozambique and South Africa [8]. Distribution of *G. brevipalpis* is limited by its habitat (forests with bush clumps for its breeding sites) and climate, specifically influenced by temperature (16 ◦C to 32 ◦C) and relative humidity (≥ 70% of relative humidity) [3,9,10].

Considering the case of Mozambique, four species of tsetse flies are found in the country, namely *G. austeni*, *G. brevipalpis*, *G. morsitans* and *G. pallidipes* [11–13]. In this country, the southern fly belt consists of two species (*G. austeni* and *G. brevipalpis*), while the central and northern belts are composed of the four tsetse species [11–13].

In the late 1890s, tsetse fly populations suffered a significant reduction, partially due to the occurrence of a rinderpest pandemic that drastically diminished the populations of both wild and domestic ungulates, particularly in southern Africa, including Mozambique [5,14]. Tsetse flies survived in small pockets allowing for a subsequent reoccurrence and expansion of the extremely reduced *Glossina* populations once the presence of game and cattle was re-established [5,14]. Nevertheless, in South Africa and Mozambique, this event was followed, between 1920 and 1970, by important vector control programs against all tsetse fly species [10,15,16]. These control campaigns reshaped the tsetse distribution to what it is today, with the dominance of *G. brevipalpis* and *G. austeni* in several zones of Southern Africa, and in particular in Mozambique [9,14–18]. In Mozambique, the southern provinces, Gaza and Inhambane, have remained tsetse-free for the last 120 years, without evidence of tsetse fly reinvasion. This separates the central population of *G. brevipalpis*, mainly in the Gorongosa National Park, from the southern population of the same species primarily situated in the Maputo National Park [8] (Fig. 1).

**Fig 1.**
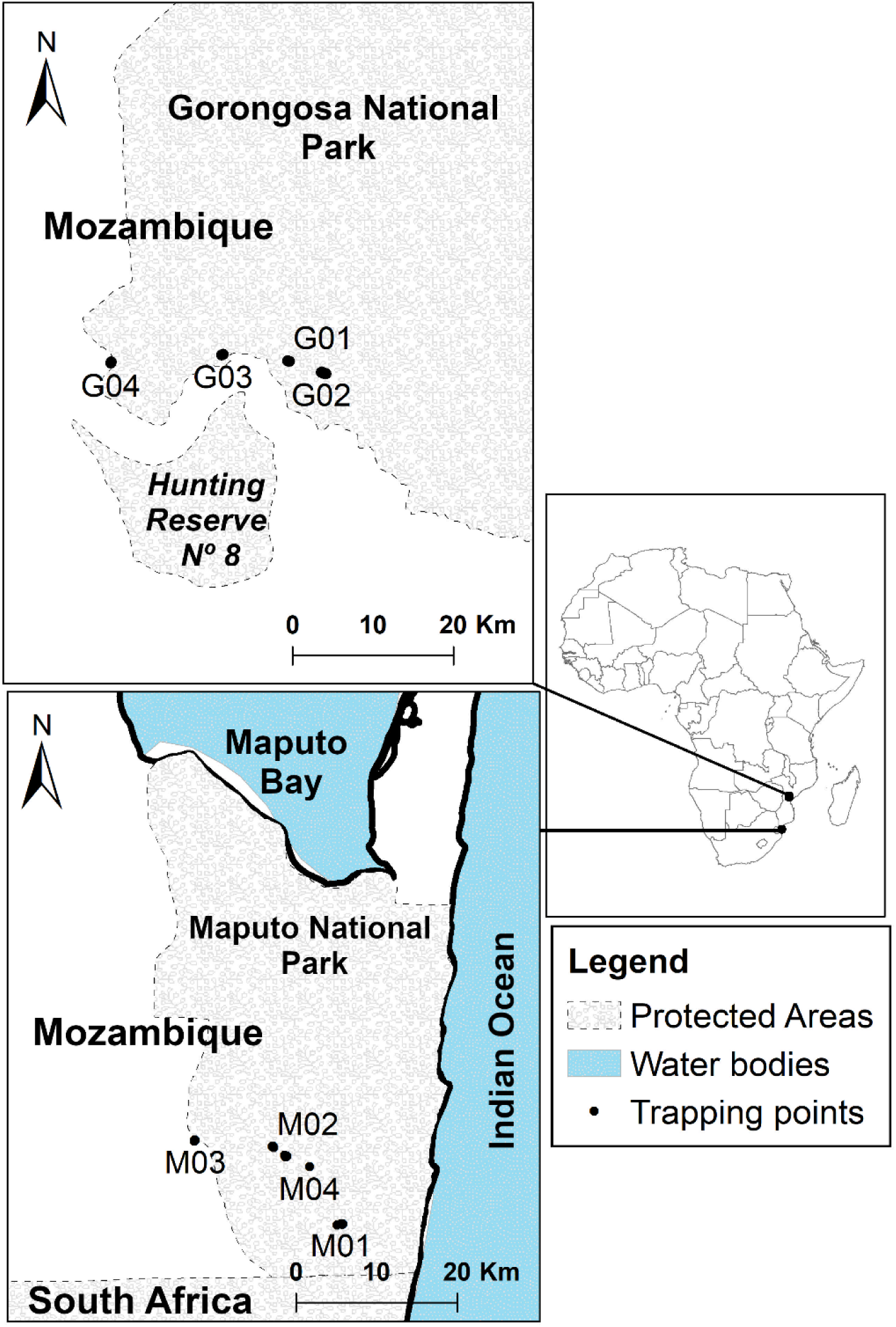
Map of sampling sites (indicated by black dots) of *Glossina brevipalpis* in both Gorongosa National Park and Maputo National Park. Traps contained in each site and their precise GPS coordinates can be seen in supplementary Table S1.

One very efficient strategy to reduce trypanosomosis incidence is vector control [19]. In order to design the most efficient control strategy, understanding the dynamics of the existing populations of *Glossina* is important [20]. It is not only essential to know the existing species and their distribution within a country, but also to understand the genetic structure and gene flow between their populations [20,21]. Microsatellites have been extensively used for tsetse fly population genetics over the last two decades [21–25].

To develop more effective and sustainable vector control strategies in Mozambique, a better understanding of gene flow between Gorongosa National Park and Maputo National Park is crucial. With this goal in mind, the present paper aims to assess the genetic diversity and gene flow between the central and southern populations of *G. brevipalpis*.

## Methods

### Study sites and sample collection

Gorongosa National Park (GNP) is located in the central region of Mozambique. It was originally established as a hunting reserve in the 1920s and now includes 3,770 km^2^ of protected land, which is composed of a mix of forest and savannah landscapes, with a high diversity of fauna and flora. This area includes four species of tsetse flies, namely *G. pallidipes, G. morsitans, G. brevipalpis* and *G. austeni*. About 840 km to the south, in the most southern point of the country, there is a tsetse hot spot that partially overlaps with the Maputo National Park (MNP). The park was established in 1960 as a national reserve to protect the extensive population of elephants found in the region. Today, it covers a total of 1,718 km^2^, comprising various ecosystems including forests that harbour two species of tsetse flies, *G. brevipalpis* and *G. austeni*. Between these two parks, including northern Maputo province, Gaza and Inhambane provinces, a putatively tsetse-free area has existed since the rinderpest pandemic of 1896 [15]. For this reason, these two parks were chosen as sampling areas to study gene flow of *G. brevipalpis* populations between the south and central regions of Mozambique.

Six hundred and seventy-nine (679) *G. brevipalpis* specimens were collected in GNP (342) and MNP (337) using 49 traps (36 H-traps [26] and 8 NGUs traps [27], enhanced with odour attractants (1-octen-3-ol and acetone) [26–28]). All flies were captured between June 14 and July 22 2022, which corresponds to less than one tsetse fly generation [7]. Twenty-nine traps were deployed in GNP and 20 in MNP (Fig 1). In each park we selected four different locations in which two to six traps were placed covering a total distance of 1 km (Fig 1). All traps were georeferenced using the eTrex 10 GPS (Garmin Ltd., USA). The overall sampling transects, i.e. the distance between the most extreme traps, reached 25 km in GNP and 14 km in MNP.

### DNA extraction and genotyping

Four hundred and two (402) *G. brevipalpis* (GNP – 202 and MNP – 200) were selected for genotyping based on the location and condition of specimens. DNA was extracted from 3 legs per individual using the chelex method [29] after which the DNA was diluted for amplification. We used 11 microsatellite primers developed by Gstöttenmayer et al., (2023) to genotype individual flies collected from both parks: Gb5, Gb28, Gb35, Gb48, Gb66, Gb70, Gb72, Gb73, Gb92, Gb158, and Gb165. To incorporate the dyes into the PCR product, microsatellite forward primers contained the M13 adapter sequence and amplification was done with M13 primers attached to one of these four dyes: VIC, NED, PET and FAM. PCRs were carried out in a 20 µL reaction with 1x PCR Buffer, 0.2 mM DNTPs, 0.08 µM forward primer, 0.1 µM reverse primer, 0.1 µM M13 primer with dye, 0.5 U Taq polymerase and 10 µL of diluted DNA. The PCR cycling conditions were as follows: 95 ^○^C for 3 mins and then 10 cycles of 95 ^○^C for 30 s, T_a_ (58/59 ^○^C) + 5 ^○^C for 30 s, 72 ^○^C for 1 min dropping 0.5 ^○^C each cycle, then 30 cycles of 95 ^○^C for 30 s, T_a_ (58/59 ^○^C) for 30 s, 72 ^○^C for 1 min, with final elongation step at 72 ^○^C for 5 min. The PCR products were pooled by four (the different dyes) and resolved on an ABI 3500XL sequencer (Thermo Fisher Scientific, USA). Allele calling was done using GeneMapper Software v6.1 [30] and the size standard GS600LIZ (Thermo Fisher Scientific, USA).

### Data analysis

Genotypic data were formatted into the appropriate file types using Create v1.37 [31]. Analyses were done using 10 autosomal loci developed by Gstöttenmayer et al., (2023) since Gb70 is sex-linked and thus was removed from our dataset.

To describe the population genetic structure of *G. brevipalpis*, we used Wright’s *F*-statistics [32]: *F*_IS_, which measures the effect of deviation from random mating within subpopulations on inbreeding, and *F*_ST_, which measures the effect of subdivision. These were respectively estimated with Weir and Cockerham’s (1984) unbiased estimators (*θ* and *f*). Their significant deviation from 0 was tested with permutations of alleles between individuals within subsamples (for *F*_IS_) and individuals between subsamples (for subdivision). In each case, the statistic used was the *F*_IS_ estimator or the natural logarithm of the maximum likelihood ratio (*G*) [34] respectively. We also tested linkage disequilibrium between each pair of loci with the *G*-based randomization test over all subsamples [35]. For each test we implemented 10,000 permutations. Estimates and testing were undertaken with Fstat v2.9.4 [36].

Wahlund effect occurs when individuals belonging to subpopulations with different allele frequencies are admixed into the same subsample. This was first used to explain heterozygote deficits as compared to Hardy-Weinberg expectations following this phenomenon [37]. Nonetheless, Wahlund effects can also minimize subdivision measures, and increase linkage disequilibrium between loci. To determine the levels at which *G. brevipalpis* from Mozambique were subdivided, we used the Wahlund effect detection technique of Goudet et al., (1994) . For this, we considered four sampling designs, depending on the level considered: trap, location, park, and all flies together (All). If a Wahlund effect occurs at one level, a significant change should be observed. We thus computed *F*_IS-trap_, *F*_IS-location_, *F*_IS-park_, and *F*_IS-All_, and compared them using the Wilcoxon signed rank test [39] for paired data using R-commander package v2.9-5 (Rcmdr) [40,41] for R v4.4.2 [42], with the alternative hypotheses: *F*_IS-trap_ < *F*_IS-location_ < *F*_IS-park_ < *F*_IS-All_. We also compared *F*_ST_’s measured at these levels (except All). For those, we undertook the same test but with a reverse alternative hypothesis. We finally compared the proportions of locus pairs in significant LD with one-sided Fisher exact tests [43] with R v4.4.2 (command fisher.test). Because of the non-independent test series undertaken for each parameter compared, we needed to adjust the obtained *p*-values with the Benjamini and Yekutieli (BY) procedure [44], with the command “p.adjust” in R v4.4.2.

#### Quality testing of the data

We assessed the quality of our loci by checking the statistical independence between each locus pair with the *G*-based randomization test at the BY level of significance. We also checked the deviations from expected genotypic frequencies with *F*_IS_ estimates and testing as described above. We also computed 95% confidence intervals with 5,000 bootstraps over loci for averages *F*_IS_ and *F*_ST_ with Fstat v2.9.4. To obtain 95 % CI of *F*_IS_ for each locus, we used 5,000 bootstraps of individuals in each subsample for each locus with Genetix v4.05.2 [45]. We computed the average of *F*_IS_ and its 95% CI for each locus obtained with Genetix v4.05.2 weighted by the number of visible genotypes and the local genetic diversity as estimated by Nei’s unbiased estimator (*H*_S_) estimated with Fstat v2.9.4. We estimated the variation of *F*_IS_ and *F*_ST_ across loci with the standard error (SE(*F*_IS_) and SE(*F*_ST_)) obtained by jackknives over loci with Fstat v2.9.4. The presence of null alleles was assessed with several criteria as described elsewhere [46,47]. In the case of null alleles, the ratio 𝑟^SE^ = SE(𝐹^IS^)/SE(𝐹^ST^) > 2, a positive correlation is expected between *F*_IS_ and *F*_ST_, and between the number of missing data (*N*_missing_) and *F*_IS_. Correlations were assessed and their significance tested with one-sided Spearman’s rank correlation using Rcmdr v2.9-5. Null allele frequencies (*p*_nulls_) were then estimated with the EM algorithm [48] using FreeNA [49]. For this, we recoded missing data as homozygotes for allele 999 as advised [49] only for loci for which missing data indeed corresponded to true null homozygotes (see Results section). Finally, we undertook the regression *F*_IS_ ∼ *p*_nulls_ with its 95% CI of bootstraps and computed its determination coefficient and the intercept, which should correspond to the value of *F*_IS_ in the absence of null alleles (*F*_IS-0_).

#### Genetic signatures of subdivision

We estimated subdivision with FreeNA and the ENA algorithm to correct for null alleles (*F*_ST-ENA_). For this, we recoded missing genotypes suspected to correspond to true null homozygotes as 999999 as recommended [49]. Microsatellite loci generally display a high level of polymorphism, leading *F*_ST_ to reflect mutation and immigration together. To correct for this excess of polymorphism, we used the *F*_ST_’ approach [50,51]. The maximum possible *F*_ST_ (*F*_ST-max_) was computed with Fstat v2.9.4 after allele recoding by RecodeData v0.1 [51], which was used to compute 𝐹^ST-ENA^′ = 𝐹^ST-ENA^/𝐹^ST max^. Meirmans and Hedrick proposed an alternative correction (*G*_ST_") [52] which is expected to be more accurate when Nei’s *G*_ST_ is not negatively correlated with *H*_S_ [53]. Nevertheless, *G*_ST_” neither allows correction for null alleles nor the estimate of 95% CIs with available programs.

We estimated the probable number of immigrant flies exchanged between the two parks as 𝑁 ^e^𝑚 = (1 ― 𝐹^ST^)/(4𝐹^ST^), assuming an infinite Island model (*n* = infinite), or 𝑁^e^𝑚 = (1 ― 𝐹^S^)/(8𝐹^ST^), assuming a two Island model (*n* = 2) [54].

We estimated effective population sizes with five methods: the heterozygote excess method (H_ex_) from De Meeûs and Noûs, (2023); the linkage disequilibrium method (LD) [56] corrected for missing data [57]; the Coancestry method (CoA) [58]; the one locus and two loci correlation method (1L2L) [59]; and the sibship frequency method (Sib) [60]. For H_ex_ we estimated *N_e_* from the published formula in a spreadsheet program from *F*_IS_ values obtained in each subsample for each locus and averaged across loci as recommended [55]. For LD and CoA, we used NeEstimator v2.1 [61]; for 1L2L we used Estim v1.2 [59]; and finally, for Sib, we used Colony [62]. We also kept the minimum (min) and maximum (max) values and averaged all estimates across methods by weighting those with the number of usable figures obtained, as recommended [63]. In case of unusable values (i.e. “Infinite”), and in order to compare effective population sizes between the two parks, we retrieved the lower limit of the 95% CI outputted by the software used to get comparable estimates.

We then used effective population size estimates to compute the number of generations of the split between GNP and MNP subpopulations, assuming total isolation, with the formula 𝑡 = ― 2 × 𝑁e × 𝑙𝑛(1 ― 𝐹ST) (e.g. Hedrick (2005), equation 9.13a, p 502 [50]). We also extracted immigration rate (*m*) from *N_e_m* estimated with *n* = 2 or *n* = infinite and used it to compute the average dispersal distances covered by tsetse flies per generation, assuming rare transportation between the two parks, as 𝛿 = 𝑚 × 𝐷geo and its 95 % CI.

In order to help the interpretation of results, we also tried to detect a genetic signature of a past bottleneck that would correspond to the rinderpest pandemic that occurred in the years 1898 – 1901, or the massive vector control that was undertaken in the years 1948 – 1970. We implemented this test with the software Bottleneck [64]. As recommended (De Meeûs et al., (2021), p108-109 [65]), a significant bottleneck signature was recognised when the Wilcoxon test outputted significant *p*-values for the infinite allele model (IAM) and the two-phase mutation model (TPM), at least. We obtained a global *p*-value across the two parks with the generalized binomial procedure [66], using *k*’ = 2 as recommended [67], with the software MultiTest v1.2 [35]. Additionally, we used the Bayesian clustering method implemented by the software STRUCTURE v2.3.4 [68], 5,000 burning period and 50,000 MCMC iterations, number of clusters from 1 to 4 and 10 replicates. We then looked for the optimal partition with the method proposed by Evanno et al.,’s (2005) [69] with the online facility StructureSelector [70]. We hoped that this approach would help elucidate the pattern of dispersal between the two parks through the detection of probable immigrants or individuals that probably descended from immigrant parents a few generations ago.

### Ethical statement

All applicable international, national, and/or institutional guidelines for the care and use of animals were followed and all procedures performed in studies involving animals were in accordance with the ethical standards of Biotechnology Centre – Eduardo Mondlane University and the practice at which the study was conducted. The sampling procedures reported herein were authorized by the respective park authorities under permit numbers: PNG/DSCi/C231/2022 (Gorongosa National Park) and 03/01/2022 (Maputo National Park).

## Results

Of the 402 tsetse flies submitted to genotyping, 396 were successfully genotyped (i.e., with more than 4 loci amplified). These 396 genotypes (196 GNP and 200 MNP) were used for further analyses and are presented in the Supplementary Table 1.

### Significant levels of population subdivision

Significance only occurred for comparisons with *F*_IS_ and LD and between All and each of the three other sampling designs (all *p*_BY_ < 0.005). This means that migration of *G. brevipalpis* is free within the sampling areas in each park, i.e. across 25 km in GNP and 14 km in MNP. Consequently, for subsequent analyses, the level of subpopulation used was the park (i.e. Gorongosa and Maputo).

### Quality testing of the data

We found five locus pairs in significant LD, one of which (Gb5 and Gb66) remained significant after BY correction (*p*_BY_ = 0.02). There was a significant heterozygote deficit: *F*_IS_ = 0.166 in 95% CI = [0.084, 0.256] (*p*-value < 0.0002). It seemed that null alleles explained these figures well. Indeed, *r*_SE_ ≈ 3, the correlation between *F*_IS_ and *F*_ST_ was positive and significant (0.6848, *p*-value = 0.0175). Nevertheless, the correlation between *N*_missing_ and *F*_IS_ was negative (*ρ* = −0.0788, *p*-value = 0.5943) because of an excess of missing genotypes at seven loci. Indeed, it can be seen from figure 2, that the correspondence of missing genotypes with homozygotes for null alleles may be true for only three loci (Gb35, Gb73, and Gb165). Consequently, for null allele frequency estimates, we recoded missing data as 999999 only for these three loci. The resulting regression *F*_IS_ ∼ *p*_nulls_ provided a coefficient of determination (*R*²) below 0. 9 (*R*² = 0.8584, *F*_IS-0_ = 0.0316). This came from two outlier loci displaying too small *F*_IS_ when compared to the corresponding null allele frequency: loci Gb73 and Gb165. We thus chose to recode missing data for these two loci as “000000” because most blanks probably corresponded to other amplification failures than null alleles. With this new dataset (S1 table), we obtained a very good adjustment (Fig 3). The model indeed explained almost 100% of *F*_IS_ variation across loci, with a negative intercept (*F*_IS-0_ = −0.0043 in 95% CI = [-0.0704, 0.059]), compatible with pangamic populations of average effective population size *N_e_*= 116 in 95% Confidence Interval (CI) = [7, Infinite] [55].

**Fig 2.**
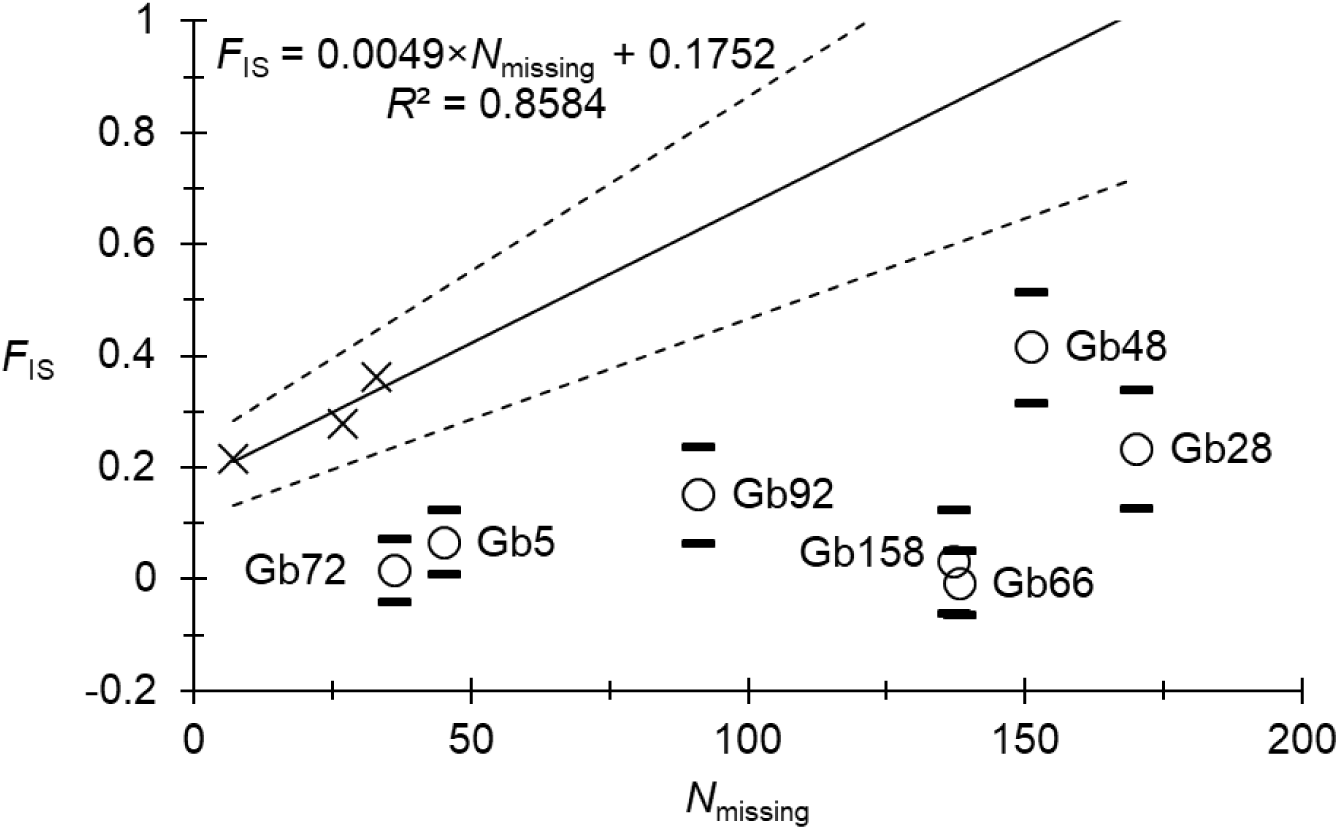
Regression *F*_IS_ ∼ *N*_missing_ for *Glossina brevipalpis* from Mozambique. The average (straight line) and its 95% CI (dotted lines) are represented. Loci for which missing data probably corresponded to null homozygotes only were used for the regression and are indicated with black crosses. Other loci are indicated by their names and empty circles with their 95% CI (black dashes).

**Fig 3.**
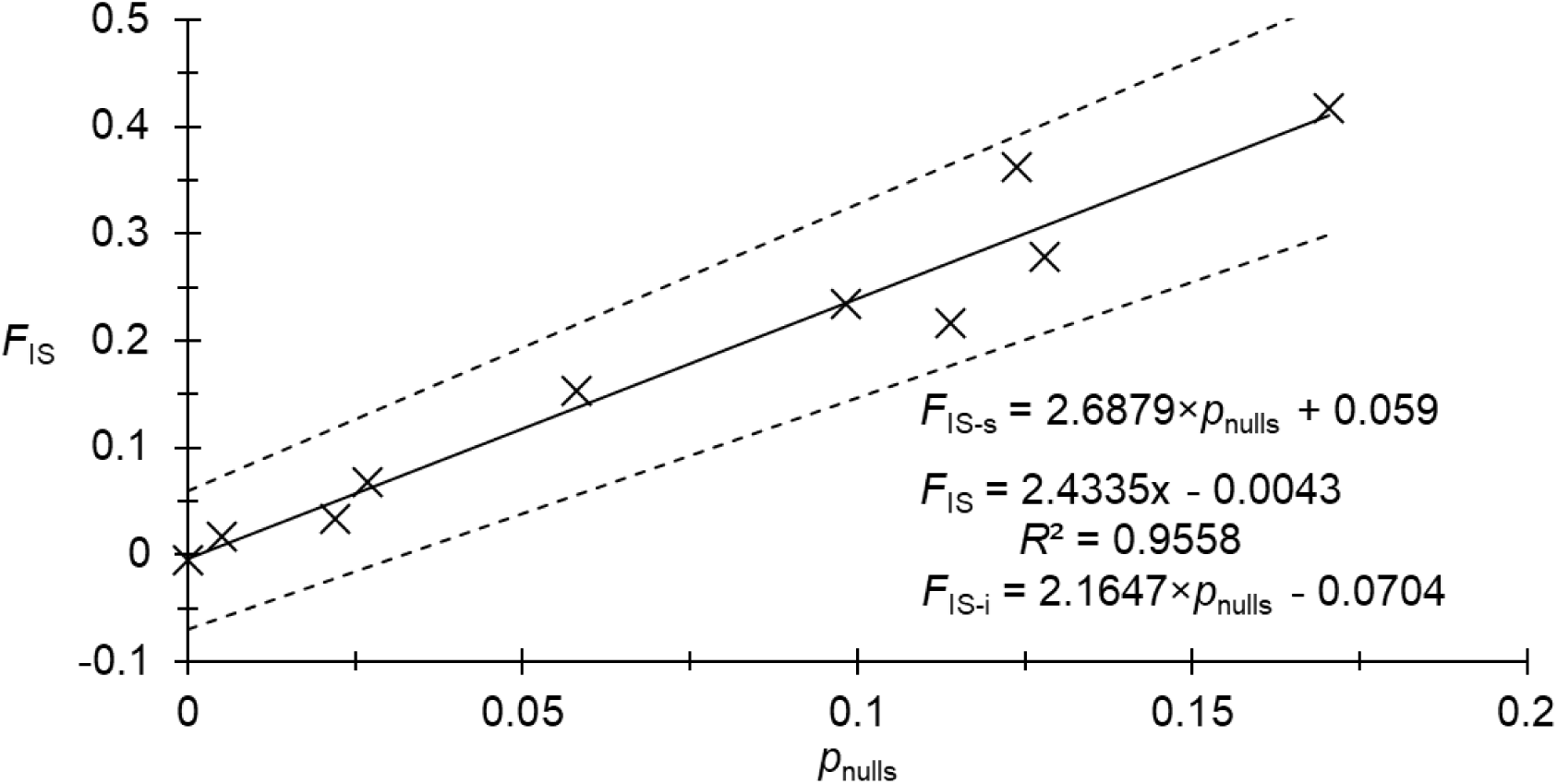
Regression *F*_IS_ ∼ *p*_nulls_ for *Glossina brevipalpis* from Mozambique obtained with 10 microsatellite loci with all missing genotypes coded as such except for locus Gb35, for which missing genotypes were considered as null homozygotes (999999) for FreeNA analyses.

To conclude, two loci, Gb5 and Gb66, are in statistical linkage and thus may introduce redundancy in multi-locus-based tests and parameter estimates. Locus Gb5 displayed a significant heterozygote deficit *F*_IS_ = 0.0674 in 95% CI = [0.0082, 0.1244], while locus Gb66 displayed a non-significant excess: *F*_IS_ = −0.0058 in 95% CI = [-0.0636, 0.0506]. A glance at https://genome.ucsc.edu/ for *G. brevipalpis* draft genome, and a BLAT [71] of primer sequences of these two loci revealed that these two loci were not found in the same scaffold. It is thus highly unlikely that these sequences are close to each other. The statistical linkage we found may therefore be a consequence of selective events driving these two markers correlatively. We thus chose to exclude loci Gb5 and Gb66 from further analyses and kept coding missing genotypes as null homozygotes (999999) for Gb35 only as regard to subdivision estimates with FreeNA.

With the eight remaining loci (Gb28, Gb35, Gb48, Gb72, Gb73, Gb92, Gb158, and Gb165), no locus pair remained significant after BY correction (all *p*_BY_ = 1). There was a significant heterozygote deficit *F*_IS_ = 0.205 in 95% CI = [0.114, 0.299] (two-sided *p*-value < 0.0002). Additionally, we observed a higher *F*_IS_ = 0.276 in MNP than in GNP (*F*_IS_ = 0.132) (*p*-value = 0.0039, Wilcoxon signed rank test [39]). This probably came from a difference in null allele frequencies, which appeared to be much higher in MNP than in GNP (0.1164 and 0.0531 respectively, *p*-value = 0.0039, Wilcoxon signed rank test [39]).

We observed a highly significant subdivision between the two parks (*p*-value < 0.0001), with *F*_ST-ENA_’ = 0.2693 in 95% CI = [0.1631, 0.3964]. With these figures, we could estimate the number of immigrants exchanged between the two subpopulations *N_e_m* = 0.68 in 95% CI = [1.28, 0.38] individuals per generation with *n* = infinite, and *N_e_m* = 0.34 in 95% CI = [0.19, 0.64] with *n* = 2.

We observed two “infinite” estimates of effective population sizes (two in MNP for Hex and CoA), which could not be taken into account for the average. Consequently, the following estimates probably corresponded to underestimations of the real average effective population size. The average effective population size was *N_e_* = 744 in minimax = [576, 912]. GNP displayed a smaller *N_e_* = 469 in minimax = [21, 2068] compared to MNP with *N_e_* = 1202 in minimax = [110, 3328], but the significance of this observation could not be tested. With the isolation hypothesis, and with these *N_e_* values, we computed that the subpopulations from the two parks needed to have split 77 years ago with a 95% CI = [42, 122], and with a minimum and maximum minimax = [32, 149]. In terms of dates, this would lead to years 1946 in 95% CI = [1900, 1980], and the oldest date going back to 1873. In case of still ongoing exchanges of immigrants, we computed that this would have required an average dispersal distance of less than 1 km per generation for *n* = 2, and between 1 and 2 km per generation for *n* = infinite.

There was no obvious signature of a bottleneck, as *p*-value = 0.0365 and *p*-value > 0.25 (maximum possible with *k* = 2) for the IAM and the TPM models.

Bayesian clustering expectedly produced an optimal partition with two clusters, but with multiple miss-assignments (Fig 4). Upon further inspection, these miss-assigned individuals seem to contain several missing genotypes. Indeed, a correlation test between the probability of belonging to the park of origin and the number of missing data in each individual fly happened to be highly significant (*ρ* = −0.1804, *p*-value = 0.0002). We thus reran STRUCTURE keeping individuals with the fewest number of missing data. Exploring data with individuals with at least less than 40% missing genotypes still provided a significant signature (*ρ* = −0.1808, *p*-value = 0.0002). With 30% missing data, the correlation was not significant anymore (*ρ* = −0.0633, *p*-value = 0.136). We thus concluded that miss-assigned individuals with more than two missing data (i.e. more than 25% of the Multilocus genotype) rather corresponded to errors, while other misplaced individuals (with a 75% complete genotype at least), probably corresponded to individuals that inherited genes from recent immigrants or even represented recently introduced individuals (if with more than 90% genomic match). This would correspond to three recently introduced individuals (all in GNP), and 51 individuals that inherited genes from more or less recent immigrants (31 in GNP and 20 in MNP). If we consider that even one missing locus was enough to bias the assignment probabilities, then we could not identify any recent immigrant.

**Fig 4.**
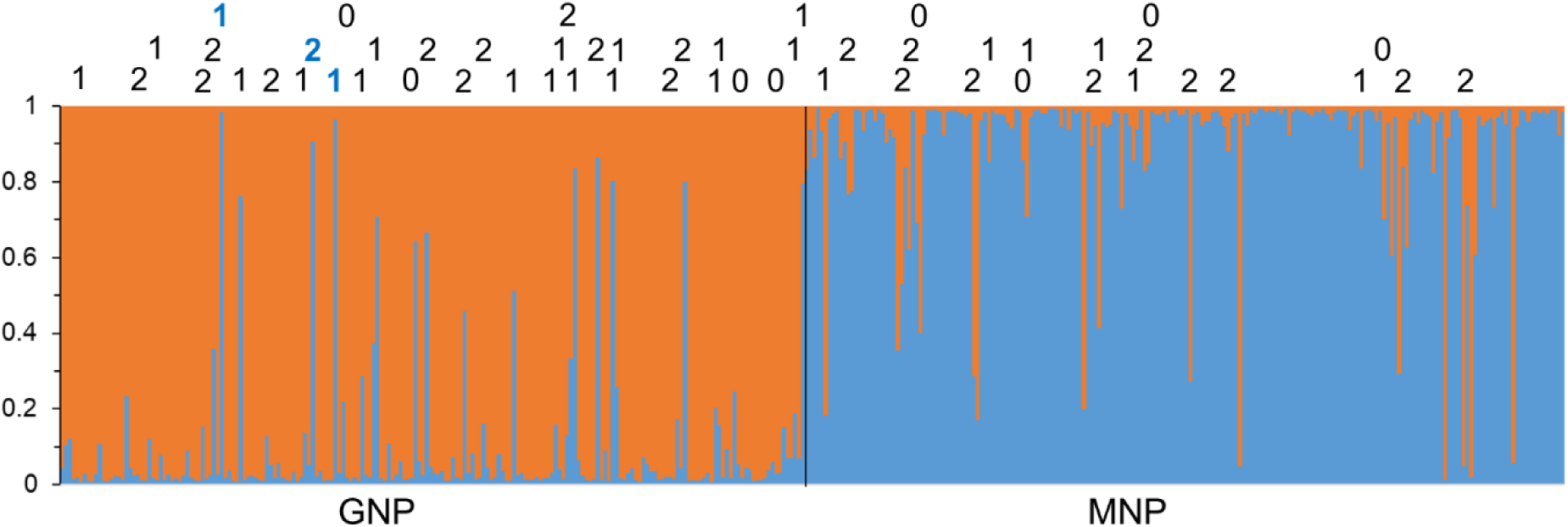
Probabilities of assignment of *Glossina brevipalpis* individuals from the two parks of Mozambique, Gorongosa National Park (GNP – orange) and Maputo National Park (MNP – blue), after Bayesian clustering and for the optimal partition obtained (K = 2). Information on the top indicate the number of missing genotypes for the individuals that were the least assigned to their park of origin, with a threshold of 0.9, and displaying less than three missing genotypes. Individuals in bold blue are more than 90% miss-assigned and assimilated to new immigrants from MNP in GNP. Numbers in black are introgressed with alien alleles inherited from some immigrant ancestors in the past.

## Discussion

During this study, we faced many genotyping problems, with a substantial number of missing genotypes at several loci. Most missing genotypes were due to PCR failures and not to null alleles, as Gb35 was the only locus for which missing data corresponded to null homozygotes. Fortunately, this did not alter most of our results as we could demonstrate that heterozygote deficits were fully explained by null alleles. We also could compute genetic differentiation accordingly. Nevertheless, during the Bayesian clustering process, such missing data significantly altered the probability of assignment of the most affected individuals to their population. Indeed, only the dataset with individuals with less than 25% of missing genotypes did not produce a significant effect of missing genotypes on probabilities of assignment.

Effective population sizes appeared substantial (*N_e_*= 744 in minimax = [576, 912]), though probably underestimated. These appeared much larger than previously reported (*N_e_* = 76 in minmax = [16, 192])[72]. Our much larger subsamples provides the most probable explanation for this discrepancy.

Subdivision between parks was highly significant. With a two-subpopulations model, this subdivision appeared compatible with the number of immigrants *N_e_m* = 0.34 in 95% CI = [0.19, 0.64] exchanged per generation. Translated in years, this would mean 2 in 95% CI = [1, 4] individuals per year, exchanged between the two parks, or a total separation dating to 1946 on average with a time window between 1900 and 1980. This would be in line with both the rinderpest pandemic (1898-1901) and the last vector control campaign, which was implemented between 1948 and 1970 in the area [5,10,15,16,73]. Nevertheless, no signature of any bottleneck could be assessed. This does not confirm a drastic decrease in tsetse populations following any of these two events. According to published data from the last 30 years [8], no *G. brevipalpis* sub-populations were reported that could link GNP and MNP in a step-by-step model. However, considering that very limited data are available from that region, this lack of reporting does not exclude the possibility that such pockets might exist and could be discovered in the future. Historical records identified a *G. brevipalpis* population pocket in Mutamba River valley near Inhambane city on the coast [74]. To this day this area has favourable conditions for *G. brevipalpis* [9], but no recent reports indicate the presence of this species. A thorough survey of these historical sites needs to be conducted. Accordingly, Bayesian clustering did not support that total isolation between the two subpopulations occurred around 1946, nor that step-by-step dispersal could take place between undocumented subpopulations connecting the two parks. Indeed, the most probable interpretation of our results is the direct exchange of (very) rare immigrants, i.e. four per year on average at most. The motorized anthropochory of these flies may be an alternative explanation for the flow of individuals between these two extremely distant populations.

Effective population size and immigration could be translated into no more than 2 km distance dispersal per generation between the two parks, which appeared much smaller than the 24 km that these flies are able to travel within a park. The difference in dispersal distance within and between the parks illustrates how low survival is for tsetse flies travelling from GNP to MNP (or back). We know that the recent introduction of wild animals occurred from one of these two parks to the other using trucks or airplanes that may have represented shelter, food and easy transport for *G. brevipalpis* [75,76]. For example, in 2019 waterbucks and oribi were transported to MNP from GNP and more recently in 2023 a pack of African wild dogs were translocated from the buffer zone of MNP to GNP [76]. This can be related to long-range dispersal distances that were recently documented in other tsetse fly species [22,77,78]. We, therefore, advise taking into consideration such a phenomenon that may account for reinvasions over very long distances.

## Conclusion

The most relevant result obtained during this work is that, despite the existence of two distant and supposedly isolated populations, there is strong evidence indicating the exchange of rare individuals. This could be due to undiscovered pocket populations between the two parks, if so a detailed entomological survey of possible habitats between the two parks needs to be conducted. A more probable explanation of our findings is that tsetse flies may have been moved between parks via motorized human transport means. Such passive dispersal should thus be investigated more thoroughly in future studies. We also recommend that a similar study be carried out on *G. austeni* populations in Mozambique that show a similar distribution pattern to that of *G. brevipalpis*. Such a study might further elucidate the tsetse dynamics between the centre and south of the country and assist in the development of efficient control strategies.

## Supplementary information

**S1 Table. Dataset of tsetse genotypes.** Three hundred and ninety-six (396) *Glossina brevipalpis* genotypes collected at Gorongosa National Park (196) and Maputo National Parks (200) in Mozambique.

## Acknowledgements

We would like to acknowledge the Biotechnology Centre of Eduardo Mondlane University and INTERTRYP IRD/CIRAD for using their laboratories and for their continued partnership throughout this study. We would also like to acknowledge Keila Zandamela and Edmilson Philimone for their assistance and dedication during this project. This project has received funding from the European Union’s Horizon 2020 research and innovation programme under grant agreement n°101000467, acronym ‘’COMBAT’’ (Controlling and Progressively Minimizing the Burden of Animal Trypanosomosis)[79].

## Authors’ contributions

All authors read, amended and/or approved the final manuscript. **Conceptualization:** Denise R. A. Brito, Fernando C. Mulandane, and Luis Neves. **Sampling:** Denise R. A. Brito, Fernando C. Mulandane, Nióbio Cossa, Hermógenes N. Mucache. **Genotyping, genotype interpretation and corrections:** Denise R. A. Brito, Adeline Ségard, Sophie Ravel. **Data analyses:** Denise R. A. Brito and Thierry de Meeûs. **Maps and design of Figures:** Denise R. A. Brito, Fernando C. Mulandane, and Thierry de Meeûs. **Writing of the original draft:** Denise R.A. Brito, Luis Neves, Thierry de Meeûs.

